# Whitening of odor representations by the wiring diagram of the olfactory bulb

**DOI:** 10.1101/515411

**Authors:** Adrian A. Wanner, Rainer W. Friedrich

## Abstract

Neuronal computations underlying higher brain functions depend on synaptic interactions among specific neurons. A mechanistic understanding of such computations requires wiring diagrams of neuronal networks. We examined how the olfactory bulb (OB) performs ‘whitening’, a fundamental computation that decorrelates activity patterns and supports their classification by memory networks. We measured odor-evoked activity in the OB of a zebrafish larva and subsequently reconstructed the complete wiring diagram by volumetric electron microscopy. The resulting functional connectome revealed an overrepresentation of multisynaptic connectivity motifs that mediate reciprocal inhibition between neurons with similar tuning. This connectivity suppressed redundant responses and was necessary and sufficient to reproduce whitening in simulations. Whitening of odor representations is therefore mediated by higher-order structure in the wiring diagram that is adapted to natural input patterns.

---

Neuronal activity patterns evoked by natural stimuli are transformed in the brain to extract relevant information. Such patterns often contain correlations and intensity variations that originate from the statistics of natural scenes and from the tuning of sensory receptors^1^. This statistical structure complicates the classification of sensory inputs because it does not usually reflect behaviorally relevant stimulus categories^2^. For example, visual scenes may be dominated by a large number of pixels representing sky while the biologically most important information is conveyed by a small subset of pixels representing specific objects (e.g., a hawk or a sparrow). Hence, correlations in sensory inputs can complicate meaningful pattern classification and object recognition. This problem can be alleviated by whitening, a fundamental transformation in signal processing that decorrelates patterns and normalizes their variance. Whitening is therefore often used early in a pattern classification process to remove undesired correlations and to optimize the use of coding space^3^.

In the visual and auditory system, whitening of individual neurons’ responses to natural stimuli supports efficient coding by redundancy reduction^4–7^. Efficient pattern classification, however, requires whitening of activity patterns across neuronal populations. This form of whitening occurs in the olfactory bulb (OB)^8–10^ where axons of olfactory sensory neurons expressing the same odorant receptor converge onto discrete glomeruli. Odors evoke distributed patterns of input activity across array of glomeruli that can overlap substantially when odorants share functional groups^11–13^. Moreover, the variance (contrast) of glomerular activity patterns varies dramatically as a function of odor concentration. As a consequence, patterns of sensory input to the OB are not well suited for concentration-invariant odor classification. The output of the OB is transmitted to higher brain areas by mitral cells, which receive sensory input from individual glomeruli and interact with other mitral cells via multisynaptic interneuron (IN) pathways (Fig. 1a). Contrary to glomerular inputs, activity patterns across mitral cells become rapidly decorrelated during the initial phase of an odor response^8,14–18^ and their variance depends only modestly on stimulus intensity^10,19^. Neuronal circuits in the OB therefore decorrelate and normalize population activity patterns, resulting in a whitening of odor representations that facilitates pattern classification. However, it remains unclear how this transformation is achieved by interactions between neurons in the OB network.

**Fig. 1.**
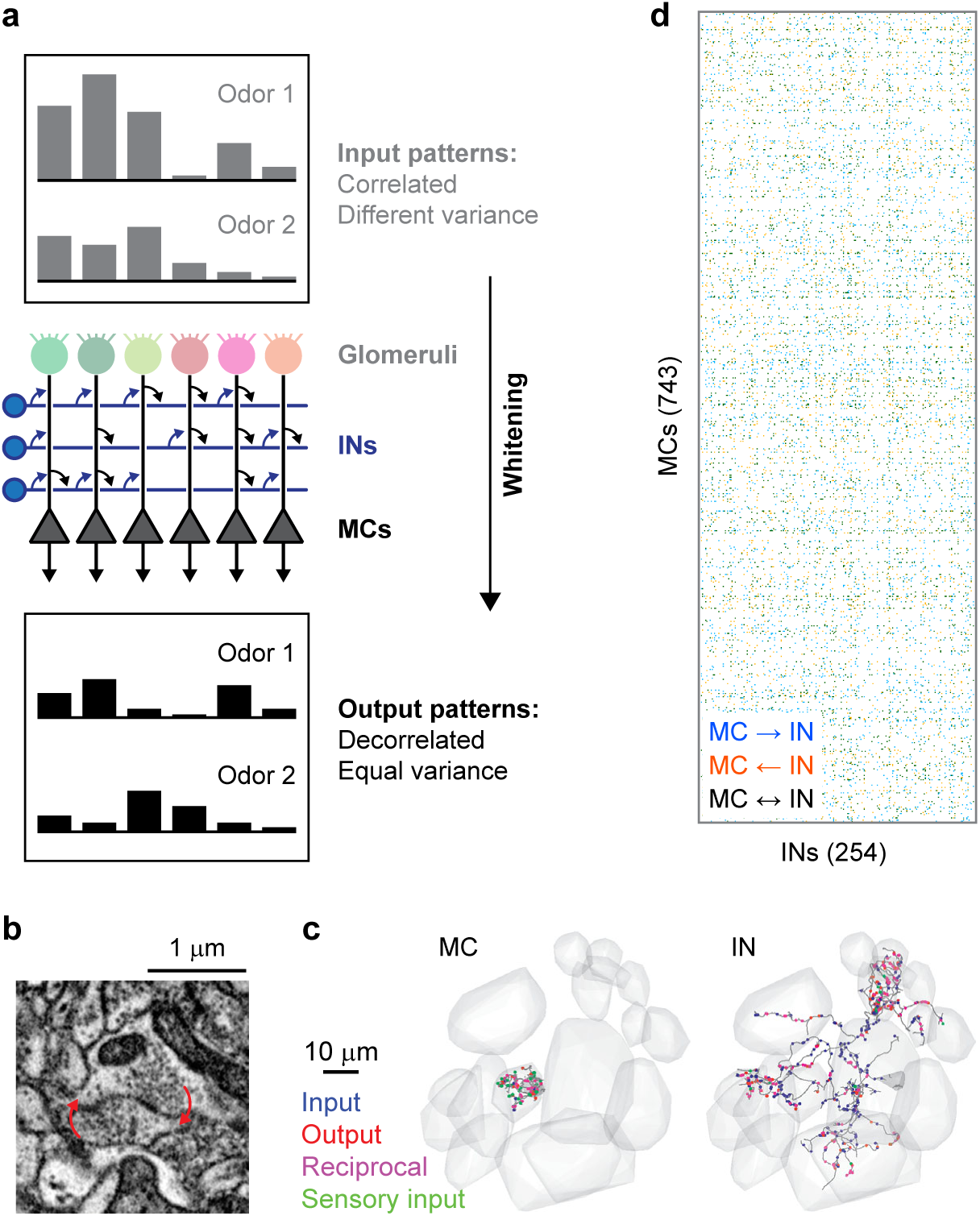
Neuronal organization and computations in the OB. **a**, Schematic illustration of whitening in the OB. Top: correlated input patterns with different variance. Bottom: decorrelated output patterns with similar variance. Center: Highly simplified illustration of the OB circuit. MCs receive excitatory input from a single glomerulus and interact via inhibitory INs. Whitening requires multisynaptic interactions between specific subsets of MCs that are mediated by INs and defined by the wiring diagram. **b**, Example of a reciprocal synapse between a MC and an IN. **c**, Reconstructions of a MC (left) and an IN (right). Gray volumes show glomeruli, dots depict synapses, colors denote synapse class (unidirectional non-sensory input [blue], unidirectional output [red], reciprocal [magenta], input from sensory neurons [green]). **d**, Simplified representation of the wiring diagram between MCs and INs (binarized connection strength). Colored matrix elements show MC→IN synapses (blue), MC←IN synapses (orange), and reciprocal synapses (black).

Efficient whitening can be achieved by transformations that are adapted to the correlation structure of input patterns^1^. Such adaptive whitening requires prior knowledge about inputs and tuning-dependent connectivity between specific cohorts of neurons. Hence, whitening of sensory representations is thought to depend on an evolutionary memory of stimulus space that is contained in the wiring diagram of neuronal circuits. This hypothesis is difficult to test in the OB because tuning and functional connectivity cannot be inferred from topographical relationships between neurons^11,20–22^. Moreover, because interactions between mitral cells are multisynaptic via INs, relevant inhibitory interactions cannot be visualized by transsynaptic tracing across a single synapse.

Adaptive whitening and other memory-based processes are likely to depend on higher-order features of neuronal connectivity that cannot be detected by sparse sampling of pairwise connectivity between individual neurons. We therefore used a “functional connectomics” approach that combines population-wide neuronal activity measurements with dense reconstructions of wiring diagrams. To achieve this goal we took advantage of the small size of the larval zebrafish brain. We first measured odor responses of neurons in the OB by multiphoton calcium imaging and subsequently reconstructed the synaptic connectivity among all neurons by serial block-face scanning electron microscopy (SBEM)^23–26^. We found that higher-order features of multisynaptic connectivity specifically suppress the activity of correlated mitral cell ensembles in a stimulus-dependent manner, resulting in a decorrelation and variance normalization. The wiring diagram of the OB is therefore adapted to the correlation structure of its inputs and mediates a whitening operation based on contrast reduction rather than contrast enhancement.

## Results

### Reconstruction of the wiring diagram and mapping of neuronal activity

We previously acquired an SBEM image stack of the OB in a zebrafish larva and reconstructed 98% of the neurons in the OB^25,26^. We now annotated the synaptic connections of all OB neurons to reconstruct the full wiring diagram. Human annotators followed each of the reconstructed skeletons and manually labeled all input and output synapses (Fig. 1b,c). Subsequently, synapses of INs were annotated a second time by different annotators. Hence, each synapse involved in MC-IN-MC connectivity motifs should have been encountered at least three times. To obtain a conservative estimate of the wiring diagram with few false positives we retained only those synapses that were annotated at least twice by independent annotators.

Each synapse was assigned a unitary weight so that the total connection strength between a pair of neurons equaled the number of synapses. The resulting wiring diagram contained 19,874 MC→IN synapses, 17,524 MC←IN synapses (Fig. 1d), and 13,610 synapses between INs. We also observed contact sites between MCs associated with the same glomerulus where plasma membranes showed strong staining but these sites usually lacked associated vesicles. We did therefore not consider synaptic connections between MCs. On average, connected pairs of MCs and INs made 3.1 MC→IN synapses and 2.9 MC←IN synapses per pair. A hallmark of synaptic connectivity in the adult OB are reciprocal dendrodendritic synaptic connections between the same MC-IN pair. In the larval OB, 52% of MC→IN synapses and 51% of MC←IN synapses were associated with a synapse of opposite direction, usually within 2.5 μm, between the same pair of neurons (Fig. 1b, bottom). Hence, reciprocal synaptic connectivity is prominent already in the larval OB of zebrafish.

Prior to preparation of the OB sample for SBEM we measured neuronal activity by multiphoton imaging of the calcium indicator GCaMP5, which was expressed under the pan-neuronal elavl3 promoter^27^. Somata observed in electron microscopy were mapped onto the light microscopy data using an iterative landmark-based affine alignment procedure followed by manual proofreading (Fig. 2a,b; Supplementary Fig. 1). Somatic calcium signals evoked by four amino acid odors (10^−4^ M) and four bile acid odors (10^−5^ M) were measured sequentially in six optical planes (Fig. 2a; Supplementary Fig. 1) and temporally deconvolved to estimate odor-evoked firing rate changes^28^. The dynamics of neuronal population activity was then represented by time series of activity vectors for each odor stimulus (232 MCs and 68 INs).

**Fig. 2.**
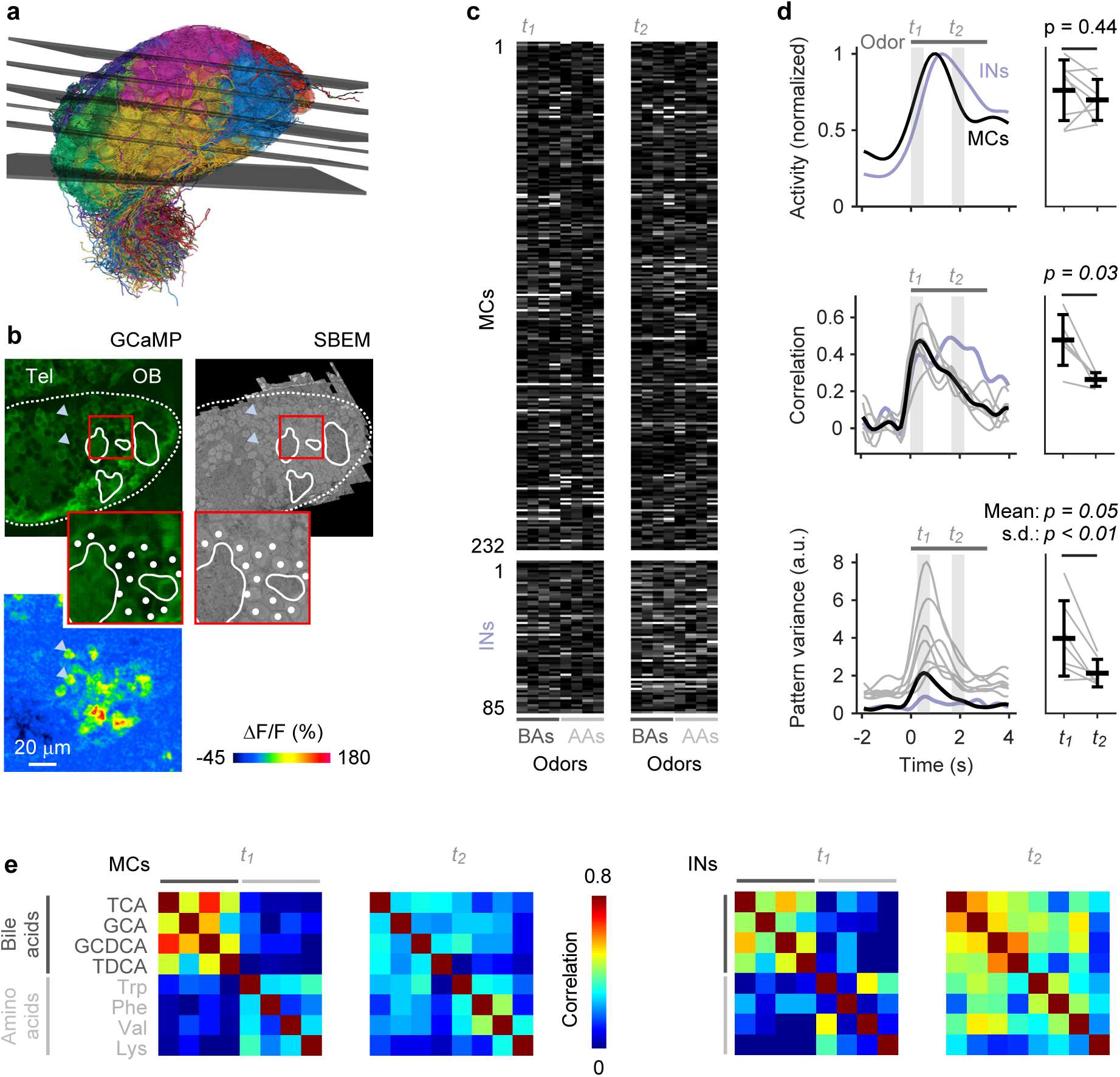
Odor-evoked population activity in the OB. **a**, Mapping of the six optical image planes selected for calcium imaging onto the EM-based reconstructions of neurons. Thickness of planes shows range of range of drift between trials. **b**, One optical image plane showing raw GCaMP5 fluorescence (left) and the corresponding oblique slice through the EM image stack (right). Dashed line outlines ipsilateral brain hemisphere; continuous white outlines show glomerular neuropil. Tel, telencephalon; OB, olfactory bulb. Region outlined by the red square is enlarged; white dots depict somata in corresponding locations. Bottom left: fluorescence change evoked by an odor stimulus in the same field of view. Arrowheads depict locations of two responsive somata in different images. **c**, Activity of MCs (n = 232) and INs (n = 68) in response to four bile acids (BAs) and four amino acids (AAs) during two time windows, *t_1_* and *t_2_*. **d**, Left: time courses of odor-evoked activity, pattern correlation (Pearson) and pattern variance. Horizontal bar indicates time of odor stimulation. Black: mean measures across MCs. Gray: individual odors (variance) or odor pairs (correlation). Light blue: mean measures across INs. Correlation was measured only between activity patterns evoked by bile acids because patterns evoked by amino acids were dissimilar already at response onset. Right: Mean measures for MCs during *t_1_* and *t_2_*. **e**, Matrices showing Pearson correlations between activity patterns across MCs (left) and INs (right) at *t_1_* and *t_2_*. Odors: TCA, taurocholic acid; GCA, glycocholic acid; GCDCA, glycochenodeoxycholic acid; TDCA, taurodeoxycholic acid; Trp, tryptophan; Phe, phenylalanine; Val, valine; Lys, lysine.

Decorrelation and contrast normalization of activity patterns across MCs have been characterized previously in the OB of adult zebrafish^8,14,15^ and mice^16–18^ where >90% of neurons are GABAergic INs. In the larval OB, in contrast, INs account for only 25% of all neurons^26^. Most of these INs are likely to be periglomerular and short axon cells because INs with the typical morphology of granule cells appear only later in development. We therefore asked whether the core circuitry present in the larval OB already performs computations related to whitening.

Correlations between activity patterns evoked by different bile acids were high after stimulus onset and decreased during the subsequent few hundred milliseconds (Fig. 2d,e). Patterns evoked by amino acids, in contrast, were less correlated throughout the odor response, which was expected because most amino acids had dissimilar side chains. To quantify pattern decorrelation we focused on activity patterns evoked by bile acids and computed the mean difference in pairwise Pearson correlations between a time window shortly after response onset (*t_1_*) and a later time window (*t_2_*). Time windows were chosen such that the mean population activity across MCs was not significantly different (Fig. 2d; p = 0.44, Wilcoxon rank-sum test). Pattern correlations across MCs, however, were significantly lower at *t_2_* than at *t_1_* (p = 0.03, Wilcoxon rank-sum test), demonstrating that MC activity patterns were reorganized and decorrelated. Activity across INs followed the mean MC activity with a small delay and did not exhibit an obvious decorrelation (Fig. 2d), consistent with observations in the adult OB^29^.

The contrast of MC activity patterns, as measured by the variance of activity across the population, increased shortly after stimulus onset and peaked slightly later than the pattern correlation. Subsequently, variance decreased and became more uniform across odors, as reflected by a significant decrease in the standard deviation of the variance across odors between *t_2_* and *t_1_* (Fig. 2d; p < 0.01, Wilcoxon rank-sum test; *t_1_* was slightly shifted relative to the time window for correlation analysis to cover the peak of the variance). Hence, MC activity patterns in the larval OB became decorrelated and contrast-normalized, consistent with the whitening of odor representations in the adult OB.

### Computational consequences of connectivity

While contrast normalization can be achieved by global scaling operations such as divisive normalization^30^, pattern decorrelation requires interactions between distinct subsets of neurons^9^. In theory, pattern decorrelation could be achieved by large networks with sparse and random connectivity^31^ but this architecture is inconsistent with the low number of INs in the larval OB. Smaller networks can decorrelate specific input patterns when their connectivity is specifically adapted to the structure of sensory inputs, suggesting that decorrelation in the OB is an input-specific transformation of odor representations that is encoded in the wiring diagram. In order to explore this hypothesis we first asked whether whitening can be reproduced by implementing the wiring diagram in a network of minimally complex single-neuron models (Fig. 3a).

**Fig. 3.**
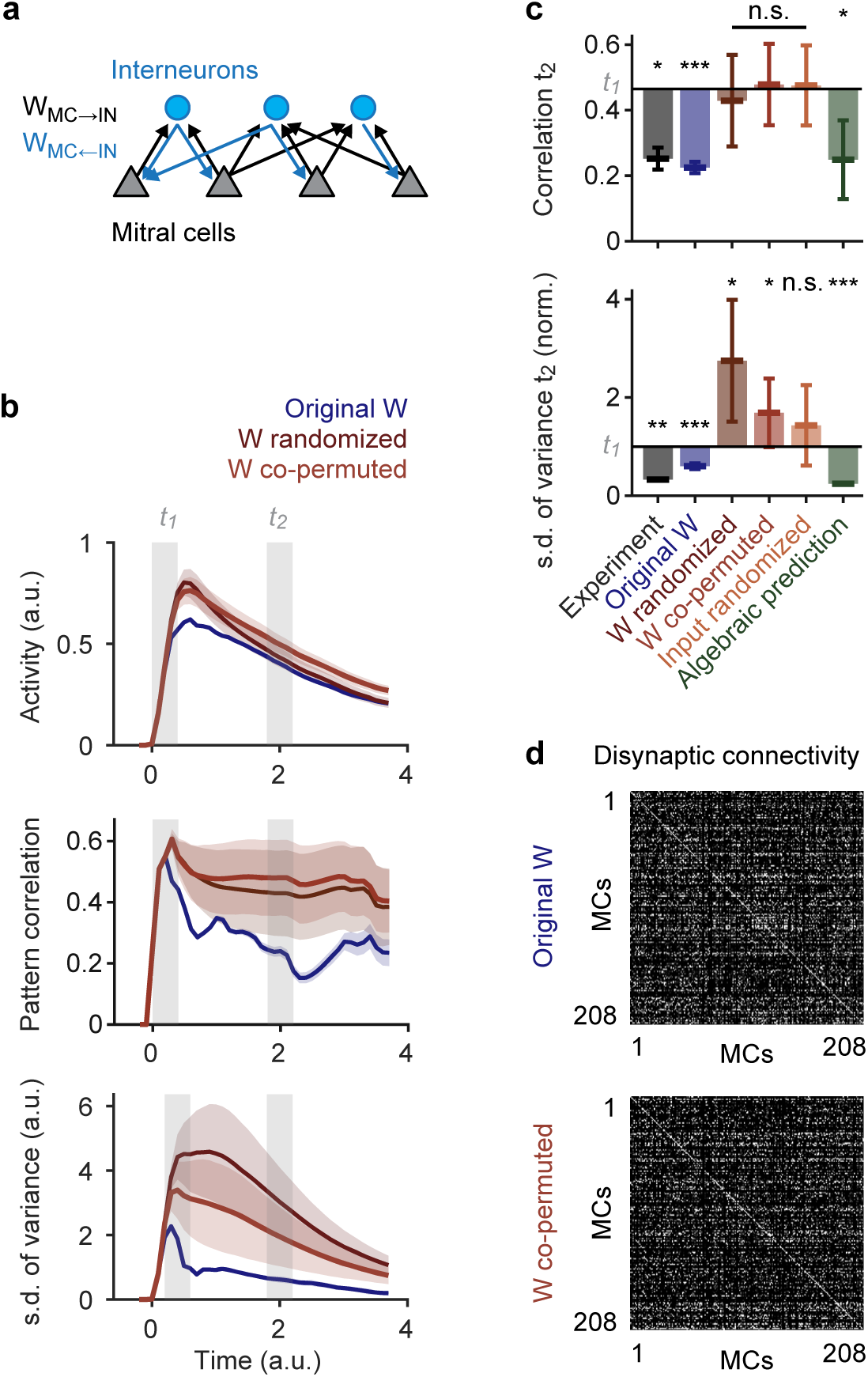
Whitening depends on connectivity. **a**, Architecture of simulated network with connections between MCs and INs. **b**, Time courses of simulated odor-evoked activity, pattern correlation and pattern variance obtained with different wiring diagrams. Blue: original wiring diagram obtained by circuit reconstruction. Dark red: fully randomized connectivity. Light red: co-permutation of feed-forward (MC→IN) and feed-back (MC←IN) connectivity. Shaded areas show s.d. across different permutations. **c**, Mean pattern correlation and s.d. of pattern variance at *t_2_*. S.d. of pattern variance is normalized to the value observed experimentally at *t_1_*. Horizontal black lines show mean experimental values at *t_1_*. Statistical comparisons of correlation and s.d. of variance were performed using a Mann-Whitney U test and an F-test, respectively. For experimental results and simulations using the reconstructed wiring diagram error bars show s.d. across odor pairs (correlation; bile acids only) or individual odors (s.d. of variance). Significance tests compare values at *t_2_* to experimental values at *t_1_*. For other simulation results, error bars show s.d. over 20 repetitions. Significance tests compare the mean over repetitions to the mean observed experimentally at *t_1_*. *, p < 0.05; **, p < 0.01; ***, p < 0.001; n.s., not significant. **d**, Top: disynaptic connectivity matrix between MCs (W_MC→IN_ * W_MC←IN_). Grayscale represents number of disynaptic MC-IN-MC connections (normalized). Bottom: example of a disynaptic connectivity matrix with the same order of MCs after co-permuting W_MC→IN_ and W_MC←IN_.

We simulated a network of threshold-linear rate neurons with 208 MCs, representing all recorded MCs with input and output synapses, and 234 INs, representing all connected INs. Connections between MCs and INs were given by the wiring diagram and excitatory sensory input into MCs was given by the odor-evoked activity pattern at *t_1_*. For simplicity, IN-IN connections were not considered. The time course of stimuli consisted of a fast initial rise followed by a slow decay^31^, approximating the response time course of olfactory sensory neurons in zebrafish^8^. Because connectivity was fixed, the final network model had only six degrees of freedom (thresholds, synaptic weight scaling factors and time constants of each neuron type).

Correlations between simulated population responses to bile acids increased rapidly and subsequently decreased. Consistent with experimental observations, the mean correlation decreased significantly between two time windows *t_1_* and *t_2_* that were chosen so that the mean activity was not significantly different (Fig. 3b,c). The variance (contrast) of activity patterns and its standard deviation across stimuli followed a similar time course but peaked slightly later than the correlation, consistent with experimental observations. Both measures decreased significantly between *t_1_* and *t_2_* (Fig. 3b,c; *t_1_* was adjusted slightly to cover the peak of the variance). Hence, a minimal network implementing the reconstructed connectivity reproduced whitening of biologically realistic inputs. When connectivity was randomized, decorrelation and contrast normalization were both abolished (Fig. 4b-d). We therefore conclude that whitening depended on the wiring diagram.

**Fig. 4.**
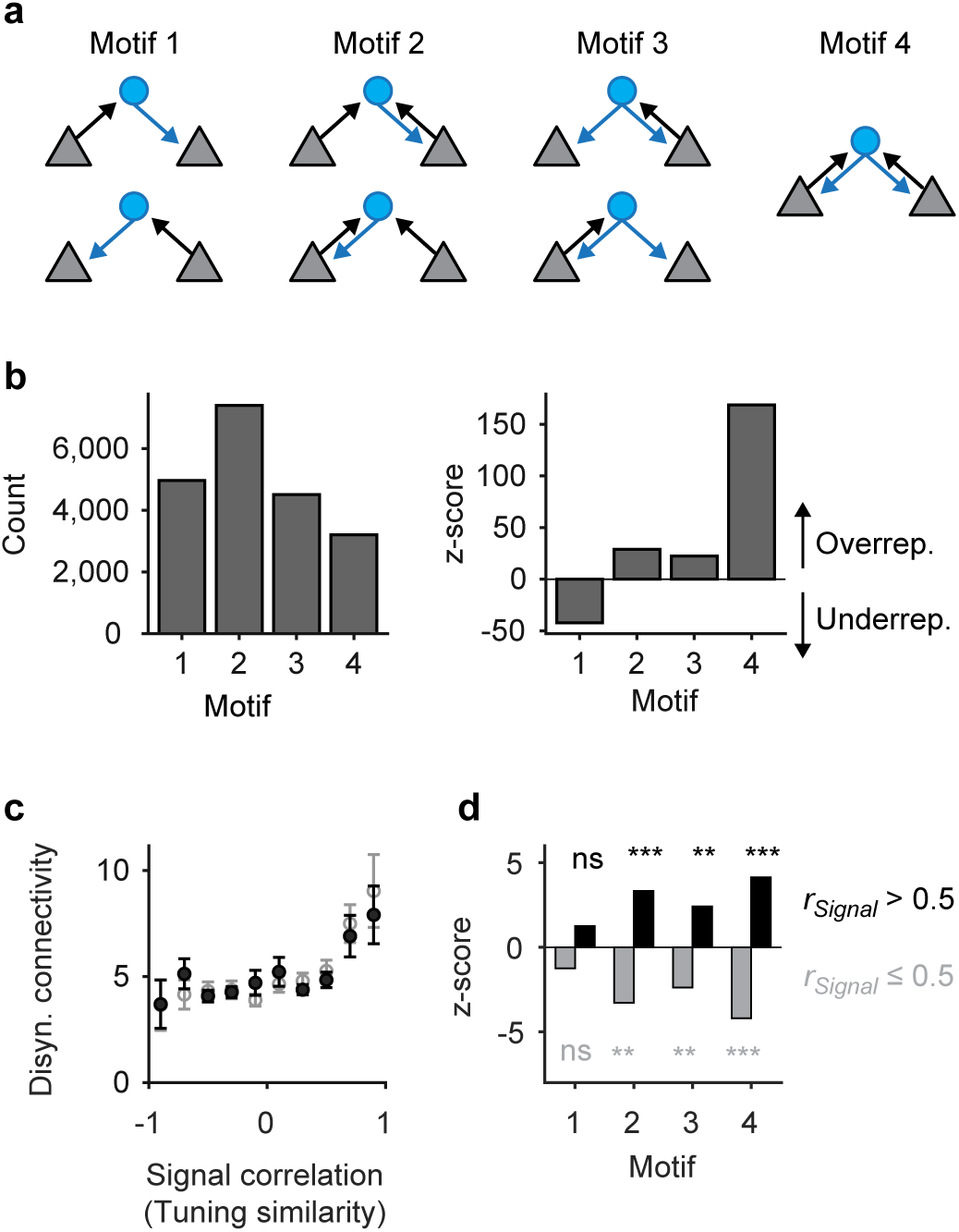
Tuning-dependent disynaptic connectivity in the OB. **a**, Classes of triplet connectivity motifs between MCs and INs. **b**, Left: number of connectivity motifs found in the wiring diagram (considering only MCs with activity measurements; n = 232). Right: z-score quantifying over-or under-representation of motifs as compared to 10,000 independent randomizations. **c**, Number of disynaptic connections between MCs as a function of tuning similarity (signal correlation; binned; mean ± s.e.m.). Black: all MCs (n = 21,528 pairs); gray: excluding MCs without at least one strong odor response (n = 7,875 pairs). **d**, Over- and under-representation of connectivity motifs among MC pairs with high signal correlation (r_Signal_ > 0.5; black) and among the remaining pairs (r_Signal_ ≤ 0.5; gray).

To further confirm this conclusion we examined whether the reorganization of activity patterns underlying whitening can be predicted from connectivity without an explicit simulation of network dynamics. Activity patterns at *t_1_* were multiplied with the feed-forward connectivity W_MC→IN_ and thresholded to generate a hypothetical pattern of IN activity. This activity pattern was then multiplied with the feed-back connectivity W_MC←IN_ to predict the pattern of feedback inhibition, which was subtracted from *t_1_*. This simple algebraic procedure reproduced both pattern decorrelation and variance normalization (Fig. 3c) but failed to do so when connectivity matrices were randomized (not shown), further supporting the conclusion that the wiring diagram contains specific information essential for whitening.

We next performed more specific manipulations to explore how whitening depends on higher-order structure in the wiring diagram. In simulations, we first applied the same permutations to the feed-forward (MC→IN) and feed-back connectivity (MC←IN). This manipulation shuffles the off-diagonal elements in the disynaptic connectivity matrix (lateral inhibition) but preserves the overall distribution of disynaptic MC→IN→MC connection strengths and the on-diagonal elements (self-inhibition; Fig. 3d). Similar to the full randomization of connectivity, this co-permutation abolished whitening (Fig. 3b,c). Moreover, whitening was abolished when input channels were permuted to produce novel input patterns with the same statistical properties and correlations (Fig. 3c). These results show that whitening is mediated by higher-order features of multisynaptic connectivity that are adapted to patterns of sensory input.

### Mechanisms of whitening

The shortest path between MCs associated with different glomeruli is a disynaptic interaction via one IN (MC-IN-MC). To identify properties of the wiring diagram that mediate whitening we therefore analyzed MC-IN-MC triplets. There are nine possible triplet configurations that represent four topological motifs (Fig. 4a). We found that the motif containing no reciprocal connection (motif 1) was underrepresented whereas the other motifs were overrepresented in comparison to randomized networks (Fig 4b). The strongest overrepresentation was observed for motif 4, which contains reciprocal connections between both MCs and the IN. Hence, MC-IN-MC triplets frequently contained reciprocal connections.

To determine whether disynaptic connectivity between MCs depends on their tuning we constructed an input tuning curve for each MC from the responses to the eight odors at *t_1_*. For all pairs of MCs we then quantified the Pearson correlation between their input tuning curves and the number of disynaptic MC-IN-MC connection paths across all motifs. The mean number of disynaptic connections increased with the input tuning correlation (Fig. 4c). Hence, triplets mediate interactions preferentially between MCs with similar tuning.

We further analyzed the relationship between triplet motifs and tuning curve similarity. Motifs with reciprocal connections (motifs 2 − 4) were significantly overrepresented among MCs with similar tuning (correlation >0.5; Fig. 4d). This overrepresentation was most pronounced for motif 4 (all connections reciprocal). Hence, disynaptic reciprocal interactions are significantly enriched between MCs with similar tuning.

In the retina, unidirectional lateral inhibition between functionally related neurons sharpens tuning curves and enhances pattern contrast^32^ (Fig. 5a, left). In idealized networks with strictly reciprocal connectivity, in contrast, inhibition does not amplify asymmetries in inputs and self-inhibition is usually larger than lateral inhibition (assuming equal synaptic strength; Fig. 5a, right). Hence, reciprocal triplet connectivity among neurons with similar tuning should primarily down-regulate, rather than sharpen, the activity of connected cohorts of neurons. The computational effects of these transformations depend on the properties of input patterns (Supplementary Fig. 2). When inputs follow overlapping Gaussian distributions, contrast enhancement can decorrelate patterns because stimulus-specific information is contained in strong neuronal responses^4,32^. However, when activity patterns overlap primarily in strongly responsive units, contrast enhancement will fail to decorrelate patterns because it emphasizes non-specific responses. In this scenario, patterns may be decorrelated by the selective inhibition of strongly active cohorts, which may be achieved by specific reciprocal inhibition (Supplementary Fig. 2).

**Fig. 5.**
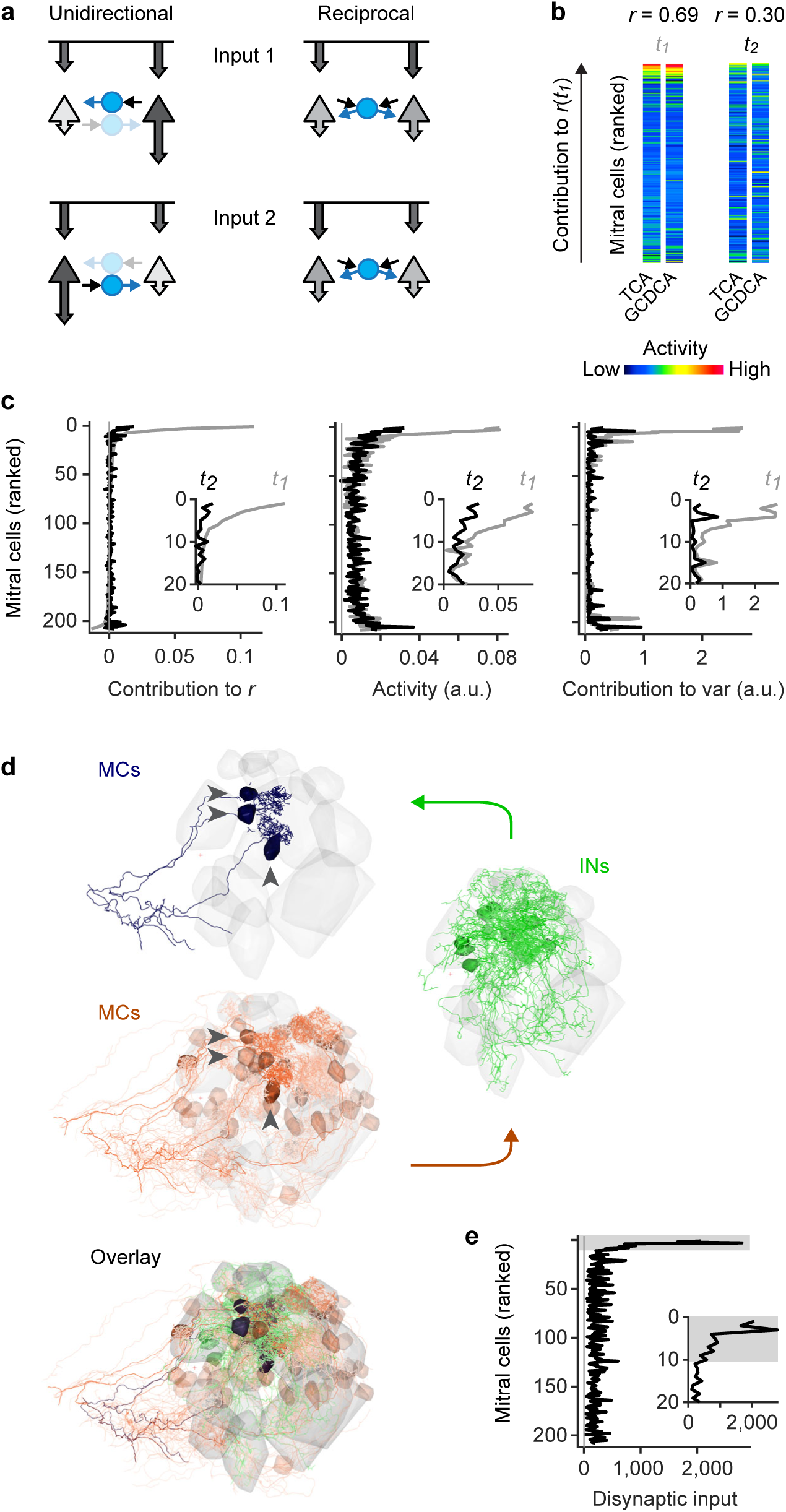
Disynaptic connectivity underlying feature suppression. **a**, Schematic illustration of contrast enhancement by unidirectional lateral inhibition (left) and down-scaling of cohort activity by reciprocal inhibition (right; feature suppression). Arrow length and grayscale indicate activity. **b**, Example of MC activity patterns evoked by two bile acids (TCA, GCDCA) that were decorrelated between *t_1_* and *t_2_*. MCs are ranked from top to bottom by their individual contribution to the pattern correlation *r* at *t_1_* (*r_i,t1_*). **c**, Left: average contribution of MCs to all pairwise correlations between activity patterns evoked by bile acids at *t_1_* and *t_2_*. MCs were ranked by *r_i,t1_* for each pair of patterns as in **b**. Sorted vectors of correlation contributions were then averaged over odor pairs. Center, right: Mean bile-acid evoked activity of MCs and mean contribution of MCs to pattern variance. MCs were sorted by *r_i,t1_* and averaged as in the left panel. Gray and black curves show correlation contribution, activity, and variance contribution at *t_1_* and *t_2_*, respectively (same sorting of individual neurons by *r_i,t1_* for all curves). Insets enlarge the top part of the curves (20 MCs with highest *r_i,t1_*). **d**, Example of disynaptic retrograde tracing of functional cohorts in the wiring diagram. Blue: three MCs with highest *r_i,t1_* for the odor pair shown in **b** (“starter MCs”). Green: 12 INs with largest number of synaptic inputs to the starter MCs. Red: 48 MCs with largest number of disynaptic inputs to the starter MCs. Transparency represents the number of synaptic connections. Note that the MCs with strong disynaptic connectivity to the starter MCs include the starter MCs themselves, consistent with pronounced reciprocal connectivity among functionally related MC cohorts. **e**, Disynaptic MC-IN-MC connectivity as a function of correlation contribution at *t_1_* (*r_i,t1_*; same ranking as in **b** and **c**). For each pair of bile acids, the 10 MCs with the highest *r_i,t1_* were selected as starter cells. Disynaptic inputs from all MCs were then represented in a vector and averaged over odor pairs. Note strong overrepresentation of disynaptic connectivity within the cohort of starter cells (gray shading).

To examine the basis of pattern correlations in the OB we analyzed population activity patterns evoked by bile acids at *t_1_*. For each pair of patterns, we quantified the contribution *r*_i,t1_ of MC *i* to the Pearson correlation *r*. Overall pattern correlations were dominated by high contributions from a small fraction of MCs. This subset of MCs was also strongly active, as observed directly when MCs were ranked by their *r*_i,t1_ (Fig. 5b,c). As a corollary, these MCs also made large contributions to the variance of neuronal activity patterns at *t_1_* (Fig. 5c). Hence, correlated odor representations overlapped primarily in strongly responsive MCs, consistent with observations in the adult OB^9^.

We then examined the changes in the activity of individual neurons underlying the decorrelation and contrast normalization between *t_1_* and *t_2_*. The activity of MCs with large *r*_i,t1_ was significantly lower at *t_2_* than at *t_1_* (Fig. 5b,c). The activity of MCs that did not strongly contribute to the initial correlation, in contrast, remained similar. As a consequence, the contribution of MCs with large *r*_i,t1_ to the overall correlation decreased, resulting in a substantial decorrelation of population activity patterns between *t_1_* and *t_2_*. Pattern decorrelation can therefore be attributed, at least in part, to the selective inhibition of MC cohorts that dominate the initial pattern correlations. MCs with high *r*_i,t1_ also made strong contributions to pattern variance at *t_1_* (Fig. 5c) because their activity was substantially higher than the population mean.

The selective inhibition of these cohorts between *t_1_* and *t_2_* changed the activity of these MCs towards the population mean and therefore decreased pattern variance and its s.d. across odors. Pattern decorrelation and contrast normalization can therefore be attributed to a common mechanism that targets inhibition to specific MC cohorts and results in contrast reduction rather than contrast enhancement.

The selective suppression of activity in cohorts of co-responsive MCs requires inhibition within cohorts to be stronger than the mean inhibition across the population. To explore how such stimulus- and ensemble-specific inhibition can arise from the connectivity between neurons we selected the 10 MCs with the highest *r*_i,t1_ for each pair of bile acid stimuli. We then determined the disynaptic MC inputs to these cohorts by retrograde tracing through the wiring diagram across two synapses. Inputs to MCs within a cohort were strongly biased towards MCs of the same cohort (Fig. 5d,e), implying that neurons in a cohort will be strongly inhibited when the cohort is activated as a whole. The specific suppression of activity underlying whitening can therefore be explained by dense reciprocal connectivity within cohorts, which suppresses the representation of stimulus features that activate a cohort.

To further explore this hypothesis we continued to analyze the mechanism of whitening in simulations. We first ranked simulated MCs by their *r*_i,t1_ for bile acid-evoked activity patterns in experiments (same ranking as in Fig. 5c). As observed experimentally, simulated MCs with large *r*_i,t1_ were strongly inhibited between *t_1_* and *t_2_* while the mean activity of other MCs remained unchanged (Fig. 6a). Simulations therefore recapitulated the mechanism of whitening in the OB and precisely predicted the underlying activity changes in individual neurons.

**Fig. 6.**
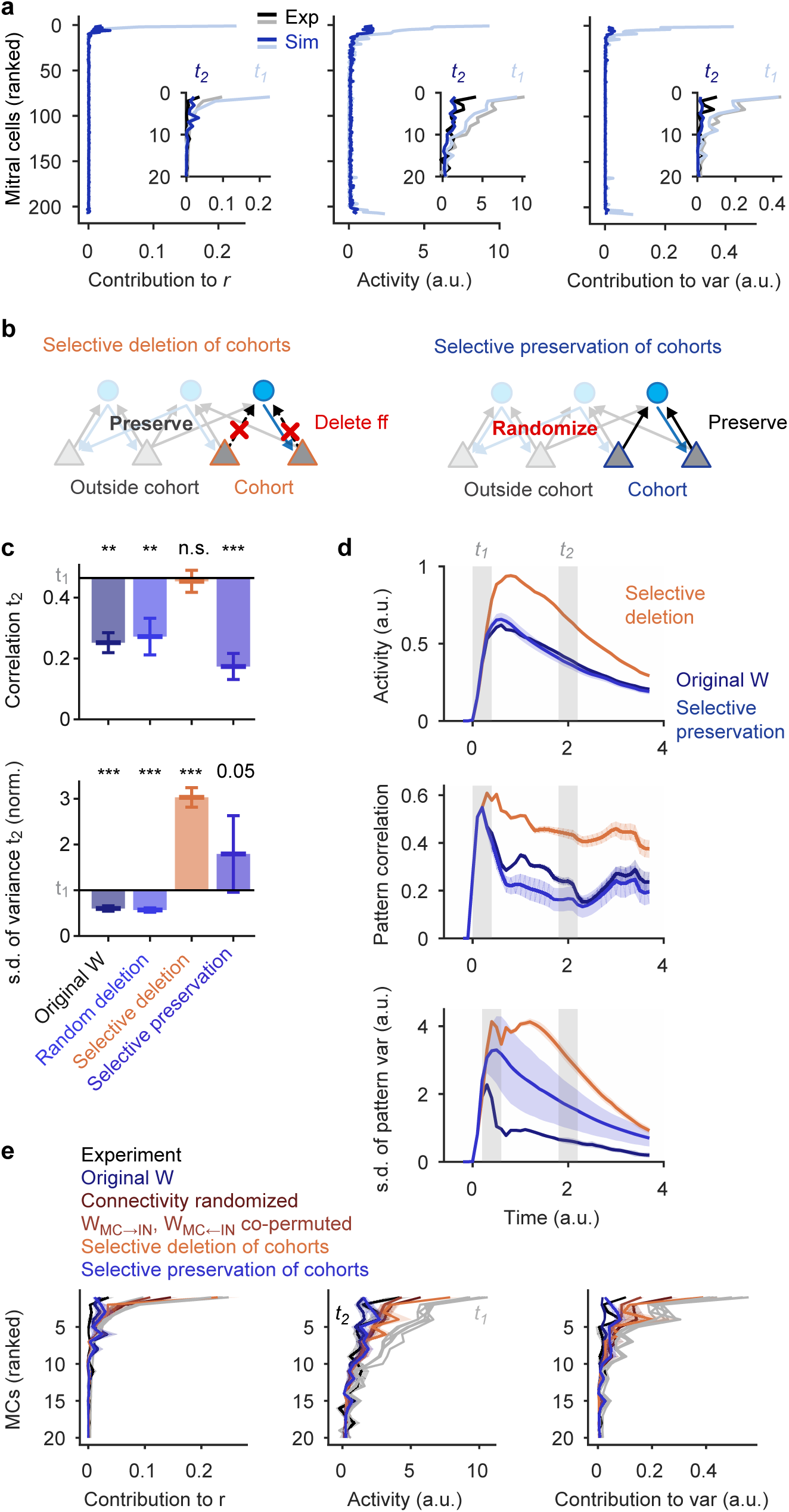
Mechanism of whitening analyzed by targeted manipulations of the wiring diagram. **a**, Mean correlation contribution, activity, and variance contribution of MCs responding to bile acids at *t_1_* (light blue) and *t_2_* (dark blue) in simulations. MCs were ranked by the correlation contribution *r_i,t1_* observed in experimental data as in Fig. 5b. Insets enlarge the top parts of the curves (20 MCs with highest *r_i,t1_*) and compare simulation results to experimental data (gray, black) for the same 20 MCs. **b**, Schematic: selective deletion and selective preservation MC cohort connectivity in simulations. **c**, Mean pattern correlation and s.d. of pattern variance (normalized) at *t_2_* observed in simulations under different conditions. S.d. of pattern variance has been normalized to the experimentally observed value at *t_1_*. Horizontal black lines show mean values at *t_1_*; vertical bars show change relative to *t_1_*. Statistical comparisons of correlation and s.d. of variance were performed using a Mann-Whitney test and an F-test, respectively. Error bars for original wiring diagram show s.d. across odor pairs (correlation; bile acids only) or individual odors (s.d. of variance); significance tests compare values at *t_2_* to experimental values at *t_1_*. Other error bars show s.d. over means from 20 simulations and significance tests compare the mean over repetitions to the mean observed experimentally at *t_1_*. **, p < 0.01; ***, p < 0.001; 0.05, p = 0.05; n.s., not significant. **d**, Time courses of pattern correlation and of the s.d. of pattern variance in simulations using different wiring diagrams. Shaded area shows s.d. across different permutations. **e**, Mean correlation contribution, activity, and variance contribution of the 20 MCs with the highest *r_i,t1_* observed experimentally and in simulations using different wiring diagrams. MCs were ranked by *r_i,t1_* observed in experimental data as in **a** and in Fig. 5c (same ranking under all conditions). Gray: *t_1_*; Colored: *t_2_*. Shading shows s.d. across 20 different permutations. Note that the reduction in correlation contribution, activity and variance contribution among MCs with high *r_i,t1_* is decreased when connectivity is modified globally or in functional cohorts, but not when connectivity of functional cohorts is preserved.

We then selected the 10 MCs with the highest *r*_i,t1_ for each pair of bile acid stimuli (19 MCs in total) and deleted their feedforward connections onto INs in the simulation (11% of all MC→IN connections; Fig. 6b, left). As a control, we deleted the same fraction of feedforward connections between random subsets of neurons. While random deletions had almost no effect, the selective disconnection of functional cohorts abolished pattern decorrelation and variance normalization (Fig. 6c,d). Ranking of MCs by their *r*_i,t1_ in experimental data demonstrated that the activity of MCs with high *r*_i,t1_ was reduced slightly between *t_1_* and *t_2_* when MC cohorts were selectively disconnected but not as effectively as under control conditions. As a consequence, these MCs continued to make large positive contributions to pattern correlation and variance at *t_2_* (Fig. 6e). These results show that the selective disconnection of functional cohorts abolished whitening because it disrupted feature suppression. We next randomized all connections except those of the 10 MCs with the highest *r*_i,t1_ for each bile acid pair (Fig. 6b, right). Results were compared to the full randomization of the wiring diagram, which reduced the inhibition of MC cohorts and abolished whitening (Fig. 3b,c). When connections of functional MC cohorts were selectively preserved, however, the inhibition of MC cohorts remained strong and pattern decorrelation was restored (Fig. 6c,d). Variance normalization was only partially rescued, presumably because preserved cohorts were selected only for their contribution to correlations between bile acid pairs and not for amino acids. The activity of MCs with high *r*_i,t1_ was strongly reduced (Fig. 6e), demonstrating that pattern decorrelation and partial variance normalization were the result of feature suppression. These results confirm that whitening is mediated by specific disynaptic interactions that suppress the activity of correlation-promoting MC cohorts.

## Discussion

We used a functional connectomics approach in a small vertebrate to explore the mechanism of whitening in the OB. Whitening is a computation related to object classification and associative memory that requires specific transformations of defined neuronal activity patterns. Such computations are thought to rely on specific wiring diagrams that are adapted to relevant inputs. Consistent with this notion, we found that whitening is achieved by specific multisynaptic interactions that cannot be described by general topographic principles or by the first-order statistics of connectivity between neuron types. Functional connectomics is therefore a promising approach to dissect distributed, memory-based computations underlying higher brain functions.

Correlations between input patterns in the OB were dominated by distinct subsets of strongly active input channels. This correlation structure is likely to reflect the co-activation of different odorant receptors by discrete functional groups^12,13^ and implies that input correlations cannot be removed efficiently by contrast enhancement^33–35^. Rather, patterns are decorrelated by the selective inhibition of strongly active, correlation-promoting MC cohorts. Pattern decorrelation is therefore achieved by contrast reduction, rather than contrast enhancement, which also supports contrast normalization.

Whitening requires specific tuning-dependent, disynaptic MC-IN-MC connectivity that may be established by molecular or activity-dependent mechanisms. Because this connectivity exists already before activity-dependent effects were detected on the morphological development of glomeruli^36^ the initial assembly of neuronal connections may rely primarily on molecular cues. Projections of INs are enriched between glomeruli that receive input from odorant receptors of the same families^26^, raising the possibility that glomerular targeting of sensory neurons^37^ and INs involve related mechanisms. However, the development of the connectivity that mediates whitening remains to be explored.

Lateral inhibition between neurons with similar tuning is often assumed to sharpen tuning curves by amplifying asymmetries in the input. In the OB, however, triplet connections between related MCs are highly enriched in reciprocal connections. This connectivity results in feedback inhibition that is independent of the precise pattern of MC input to a cohort (Fig. 5a, right) and down-scales the activity of neuronal cohorts without amplifying asymmetries in the input. Reciprocally connected MC↔IN↔MC cohorts therefore mediate feature suppression: in the presence of a feature that effectively activates a cohort, the inhibitory feedback gain within the cohort will be larger than the mean feedback gain and suppress the representation of the feature. This mechanism can explain the selective and odor-dependent inhibition of correlation-promoting MC cohorts.

Functional connectomics permitted us to test the significance of this mechanism by implementing the wiring diagram in a network of minimally complex model neurons. Simulations demonstrated that higher-order features of connectivity were necessary and sufficient to produce whitening. Precisely targeted manipulations confirmed that whitening was the result of feature suppression by reciprocal MC↔IN↔MC connectivity among correlation-promoting MC cohorts. Whitening in the OB is therefore produced by a network mechanism that differs from canonical computations in the retina and other sensory systems, presumably because the statistics of sensory inputs differ between sensory modalities.

In visual cortex, functionally related principal neurons make stronger excitatory connections than random subsets of neurons^38^. Such connectivity can arise from Hebbian plasticity mechanisms, enhance representations of sensory features, and amplify specific inputs in memory networks after learning. The disynaptic connectivity observed in the OB, in contrast, results in inhibitory interactions between functionally related principal neurons. Such connectivity cannot be achieved by monosynaptic connectivity between MCs because inhibitory synapses between MCs would violate Dale’s law. Functional connectivity in the OB is therefore similar in structure, but opposite in sign, to excitatory connectivity motifs in visual cortex. As a consequence, the connectivity in the OB suppresses, rather than amplifies, specific features in the input. Such a mechanism appears useful to attenuate the impact of irrelevant sensory inputs and to reduce undesired correlations. The mechanism of whitening by feature suppression is consistent with networks that have been optimized for whitening in a theoretical framework with biologically plausible constraints^39,40^. Hence, the mechanism of whitening observed in the OB may represent a general computational strategy in the brain.

## Acknowledgements

We thank B. Hu, A. Lüthi, P. Rupprecht and N. Temiz for comments on the manuscript and the Friedrich group for valuable discussions. C. Genoud made outstanding contributions to the acquisition of electron microscopy data. This work was supported by the Novartis Research Foundation, the Human Frontiers Science Program (HFSP; rgp0015/2010), and the Swiss National Science Foundation (SNF; CRSII3_130470/1, 310030B_152833).

## Author contributions

A.A.W. participated in all tasks. He analyzed image data, annotated synapses, supervised human annotators, analyzed data, and wrote the manuscript. R.W.F. analyzed data and wrote the manuscript.

## Data availability

EM data are available under http://doi.org/10.7281/T1MS3QN7. Other data are available from the corresponding author upon request.

---

**Supplementary Fig. 1.**
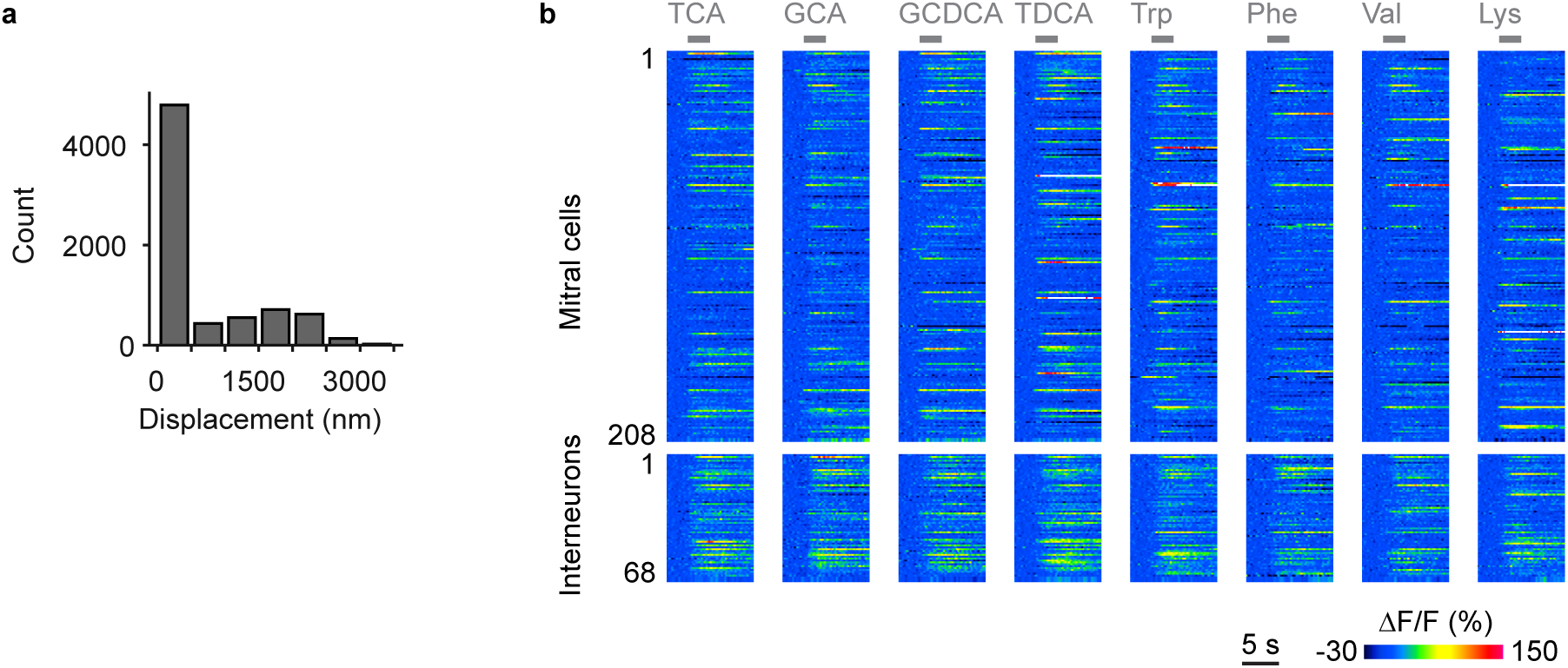
Mapping of datasets and activity measurements. **a**, Displacement of regions of interest (ROIs) during manual proofreading. ROIs representing somata were mapped from the EM dataset to optical image planes in each trial by an affine transformation that was determined by an iterative landmark-based procedure (Methods). Subsequently, the position of each ROI was adjusted manually on the optical image (n = 7,280 ROIs; six image planes with 11 trials each). The mean displacement (± s.d.) during manual adjustment (proofreading) was small (593 ± 833 nm), implying that automated mapping was highly reliable. **b**, Raw calcium signals (ΔF/F) evoked by eight odors in neurons that were present in all trials (208 MCs and 68 INs). Gray bars indicate odor stimulation.

**Supplementary Fig. 2.**
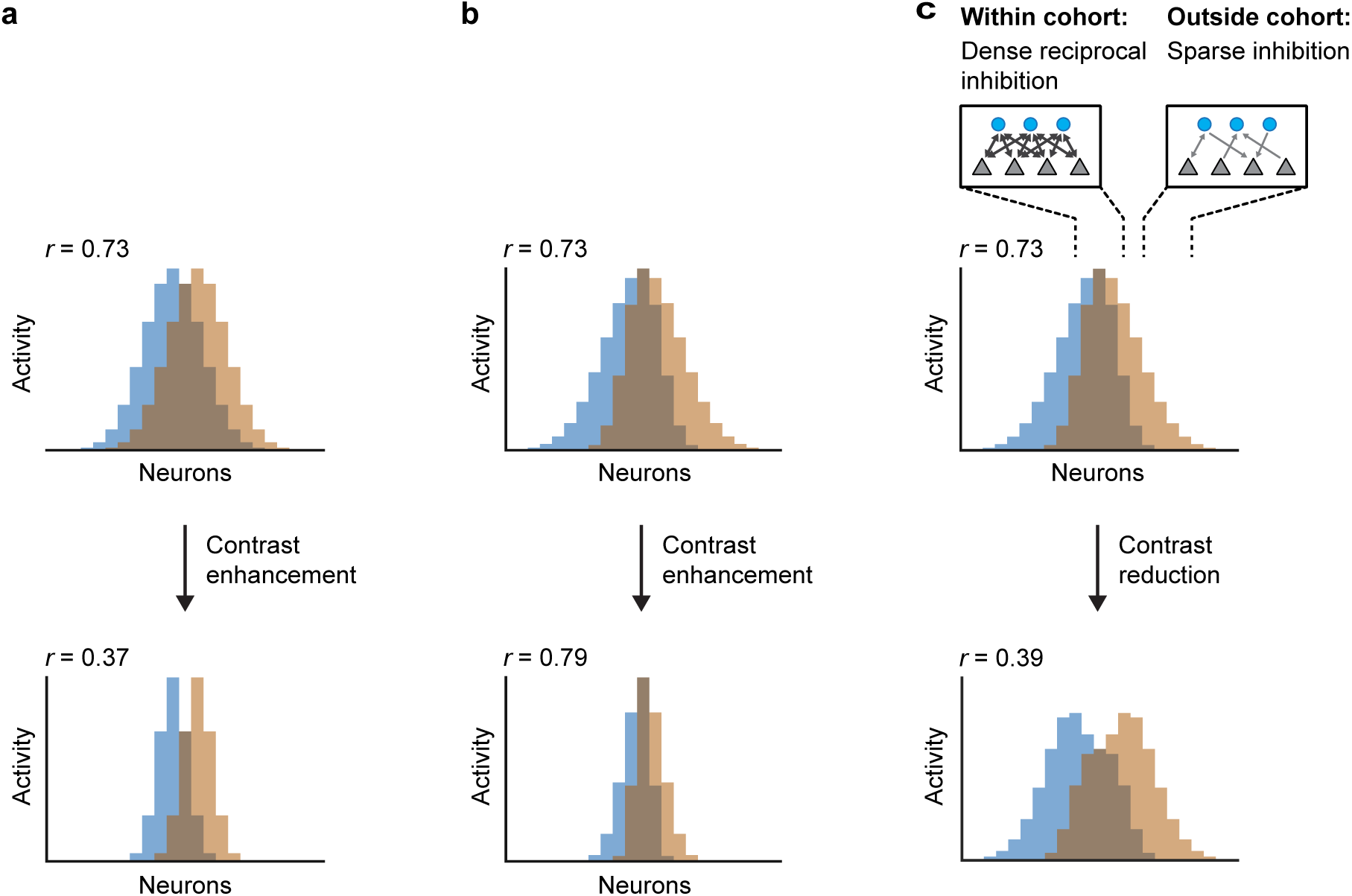
Effects of pattern transformations on pattern correlation. **a**, Effect of contrast enhancement on the correlation between displaced Gaussian patterns. In such patterns, strongly active units convey stimulus-specific information while weakly active units tend to be non-specific. Contrast enhancement therefore decorrelates patterns because it emphasizes strongly active units and suppresses weakly active units. **b**, Effect of contrast enhancement on the correlation between activity pattern that overlap in strongly active units. Contrast enhancement fails to decorrelate patterns because pattern-specific information is conveyed by moderately or weakly active units. **c**, Patterns that overlap in strongly active units are decorrelated by selective inhibition of strongly active units, which results in contrast reduction. Patterns are decorrelated because the relative contribution of moderately or weakly active units is enhanced. Selective inhibition of strongly active units is generated by dense reciprocal inhibition within cohorts of co-tuned neurons. Inhibitory feedback gain is therefore higher than the average inhibitory feedback gain within a co-tuned cohort when the stimulus feature that activates the cohort is present (feature suppression).

## Methods

### Animals and preparation

Adult zebrafish (Danio rerio) were maintained and bred under standard conditions at 26.5°C. Embryos and larvae of a double-transgenic line (elavl3:GCaMP5 x vglut:DsRed)^41,42^ in nacre background were raised at 28.5°C in standard E3 medium^43^.

Imaging experiments were performed as described previously^44^. In brief, larvae 4 - 5 days post fertilization (dpf) were contained in a small drop of aerated E3 without methylene blue or N-phenylthiourea. Larvae were then paralyzed by addition of 20 µl of fresh mivacurium chloride (Mivacron, GlaxoSmithKline, Munich, Germany)^45^ and embedded in 2% low-melting agarose (type VII; Sigma, St Louis, MO, USA) in a perfusion chamber that was inclined by 30° to improve dorsal optical access to the OBs. Agarose covering the noses was carefully removed. A constant stream of E3 (2 ml/min) was delivered through a tube in front of the nose and removed by continuous suction. Throughout the experiment it was ensured that larvae showed normal heartbeat. Larvae that were not fixed for EM recovered from paralysis after a few hours and continued to develop without obvious defects. All animal procedures were performed in accordance with official animal care guidelines and approved by the Veterinary Department of the Canton of Basel-Stadt (Switzerland).

### Odor stimulation

Odor application was performed as described^44^. In brief, odors were delivered to the nose through the E3 medium using a computer-controlled, pneumatically actuated HPLC injection valve (Rheodyne, Rohnert Park, CA, USA). All experiments were carried out at room temperature (~22°C). The odor set comprised one food odor^46^, four bile acids (glycochenodeoxycholic acid [GCDCA], taurocholic acid [TCA], taurodeoxycholic acid [TDCA] and glycocholic acid [GCA]; Sigma Aldrich, Munich, Germany) and four amino acids (Trp, Lys, Phe, and Val; Fluka, Neu-Ulm, Germany). Stock solutions of GCDCA, TCA, TDCA, Trp, Lys, Phe and Val at 5 × 10^−3^ M in E3 were kept refrigerated and diluted 1:500 (GCDCA, TCA, TDCA) or 1 : 50 (Trp, Lys, Phe, Val) in aerated E3 medium immediately before the experiment. A stock solution of GCA was prepared in 50% ethanol/50% E3 at 2.5×10^−3^ M, refrigerated, and diluted 1:250 immediately before the experiment. In a given trial, an odor was applied twice for a duration of ~3 s with an inter-stimulus interval of 60 s. Successive trials with different odors were separated by at least 2 min.

### Multiphoton calcium imaging

Multiphoton imaging was performed using a microscope equipped with a mode-locked Ti:sapphire laser (SpectraPhysics) and a 20× objective (NA 1.0, Zeiss) as described^47^. GCaMP5 was excited at 910 nm and emission was detected through green (535 ± 25 nm) and red (610 ± 37.5 nm) emission filters in separate channels. Images (256 × 256 pixels) were acquired at 128 ms per frame using SCANIMAGE and EPHUS software^48,49^ for a total of 2 min in each trial. Trials were performed sequentially in six focal planes that were separated by approximately 10 µm along the dorso-ventral axis of the OB. The field of view covered the entire cross-section of the OB and parts of the adjacent telencephalon. Ten stimulus trials (nine odors and one E3 control), each including two odor applications, were performed in each focal plane. The order of stimuli was E3, food, GCDCA, TCA, TDCA, GCA, Trp, Lys, Phe, Val. In addition, 2 min of spontaneous activity were recorded in each focal plane. After completion of all trials a stack of images covering the whole olfactory bulb was acquired with a z-step interval of 0.5 µm.

### Automated drift correction

Slow mechanical drift, which may be caused by capillary forces acting on the agarose matrix^50^, was corrected for by an automated routine. This routine acquired a small stack (± 3 µm around the focus; 0.5 µm steps) and compared images to a reference by cross-correlation after standardizing image columns and rows. The field of view was then automatically translated in X,Y and Z to maximize the cross-correlation to the reference.

### Electron microscopy

Preparation and imaging of this sample have been described previously (Wanner et al. 2016a, Wanner et al. 2016b). Briefly, tissue was stained *en bloc* with osmium, uranyl acetate and lead aspartate using an established protocol^51,52^ with minor modifications and embedded in Epon resin with silver particles to minimize charging^25,26^. Multi-tile images were acquired in high vacuum using a scanning electron microscope (QuantaFEG 200; FEI) equipped with an automated ultramicrotome inside the vacuum chamber (3View; Gatan). Section thickness was 25 nm, pixel size was 9.25 × 9.25 nm^2^, and the electron dose was 17.5 e^−^nm^−2^. The dataset comprised 4,746 successive sections of which one section was lost due to technical problems. The final stack was cropped to a size of 72.2 × 107.8 × 118.6 μm^3^.

### Neuron reconstruction and synapse annotation

Skeletons of all neurons in the OB were reconstructed previously as described^25,26^. Briefly, three independent skeletons of each neuron were generated manually from seed points at somata. Skeletons were converged and mismatches were corrected as described, and high accuracy was verified by measures of precision and recall^26^. Tracing was performed using KNOSSOS (www.knossostool.org) or PyKNOSSOS (https://github.com/adwanner/PyKNOSSOS). Most skeletons were generated by a professional high-throughput image annotation service (www.ariadne.ai).

Synapses were annotated manually using PyKNOSSOS in “flight” mode^25^. In the default configuration, PyKNOSSOS displays image data in four viewports: the YX viewport (imaging plane) and three mutually orthogonal viewports of arbitrary orientation. In “flight” mode, the latter is perpendicular to the direction of the current neurite. We found that this “auto-orthogonal” view increases tracing speed and facilitates the identification of branch points and synapses. Annotators followed skeletonized reference neurons along pre-calculated paths to ensure that all neurites were annotated. Most synapses were annotated by a professional image annotation service (www.ariadne.ai).

Synapses were identified by a cloud of vesicles that touched the plasma membrane, often at a site of intense staining. Annotators defined synapses by placing three nodes: (1) a node in the presynapse, (2) a node in the synaptic cleft, and (3) a node in the postsynapse. Nodes in the presynapse and postsynapse are skeleton nodes of the pre- and postsynaptic neurons if these skeletons are available. In addition, annotators assigned a confidence level c to each synapse. This confidence level was introduced because synapse identification is not unambiguous; rather, human experts can disagree whether a given structure is a synapse or not even when image quality is high.

Synapses were then classified as either “input synapse”, “output synapse”, “sensory synapse” or “unknown”. Input and output synapses are synapses of the reference neuron with the corresponding directions, excluding synapses with sensory neurons. Sensory synapses are input synapses received by the reference neuron from axons of sensory neurons, which were identified by their dark cytoplasm^53^. Unknown structures resemble synapses but do not display all characteristic features. These structures often included an intense staining of the membrane but no clearly associated vesicle cloud. We therefore speculate that some of these structures may be gap junctions.

We first annotated input and output synapses of all MCs and INs independently of each other. Hence, each synapse should have been encountered twice, once from the presynaptic and once from the postsynaptic side. Synapses of INs were then annotated again by different individuals, resulting in a 3-fold redundancy for each MC-IN synapse. In order to minimize the number of false positives the final wiring diagram retained only those MC-IN synapses that were annotated on the MC and at least once on the IN.

Each synapse was assigned a unitary weight. As a consequence, the strength of the connection between two neurons in each direction was given by the number of synapses between this pair of neurons. In addition, we tested two other methods to determine synaptic strength. First, connection strength was binarized such that all connections had strengths 0 or 1, independent of the number of synapses. Second, we defined the weight of a synapse as its mean confidence level c, and the total weight of a connection as the sum of the confidence levels of all synapses. In addition, we tested various confidence thresholds to discard synapses with low confidence before determining the weights. Similar results were obtained with all methods and a wide range of confidence thresholds, implying that results are highly robust.

### Correlation between multiphoton and SBEM image stacks

Mapping of multiphoton to SBEM image data may be complicated by (1) mechanical distortions introduced by the sample preparation procedure, (2) shrinkage due to loss of extracellular space induced by chemical fixation^54^, and (3) developmental changes occurring during the approximately three hours between the first calcium imaging trial and the final fixation of the tissue. Initial observations indicated that distortions between image datasets were mostly linear (rotation, translation, shrinkage) while non-linear distortions appeared minimal and developmental changes were negligible. We therefore used an affine transformation to map multiphoton images into the SBEM stack, followed by manual fine adjustment of regions of interest (ROIs) for the extraction of calcium signals.

An initial affine transformation matrix was fitted to a set of corresponding points that were selected manually in both datasets. The EM volume was then transformed onto the two-photon images, the position of existing points were optimized manually, and additional pairs of corresponding points were selected. The transform was then re-calculated based on the updated set of landmarks and this procedure was iterated until asymptotic behavior was observed.

All somata of the OB were outlined manually in the SBEM dataset and mapped onto the time-averaged multiphoton fluorescence images of each trial, resulting in 7280 mappings of somatic outlines in the SBEM dataset to regions of interest (ROIs) in 66 multiphoton images (11 trials at each of six optical planes). The position of all ROIs was then manually adjusted to optimize the mapping in each trial. The average displacement of ROIs during manual adjustment was small (593 ± 833 nm; mean ± s.d.; Supplementary Fig. 1), demonstrating that the accuracy of the initial affine mapping was already high.

### Analysis of calcium signals

Individual frames of multiphoton image time series were low-pass spatially filtered with a mild 2D Gaussian kernel (σ = 1.2 pixels). Baseline fluorescence F was calculated as the average fluorescence during a 2 s window before response onset. Traces representing relative changes in fluorescence (ΔF/F) in each ROI were averaged over the two successive odor applications in each trial and band-pass filtered in time using a Butterworth filter with a cutoff frequency of 0.2 times the frame rate. The average population response onset (t = 0) was determined manually from all raw ΔF/F traces and fixed for all trials. Firing rate changes of neurons represented by individual ROIs were estimated by temporal deconvolution of calcium signals as described^28^ using standard parameters (τ_decay_ = 3 s, *thr*_noise_ = 0).

Analyses of population activity were restricted to neurons represented by ROIs with a radius ≥2 pixels in all trials (corresponding to an area of 3.14 µm^2^; 232 MCs and 68 INs). For network simulations and mechanistic analyses of whitening we considered only the 208 MCs that were pre- and post-synaptic to at least one IN and excluded 24 presumably premature MCs. Population responses to different odors were compared by calculating the Pearson correlation coefficient between the population activity vectors of MCs for the different stimuli at a given time point after response onset.

### Network modeling

Excitatory MCs and inhibitory INs were simulated as threshold-linear units with a state variable representing firing rate. The *r*^*i*^(*t*) and *u*^*j*^(*t*) representing firing rates of MC *i* and IN *j*, respectively, followed the equations of motion

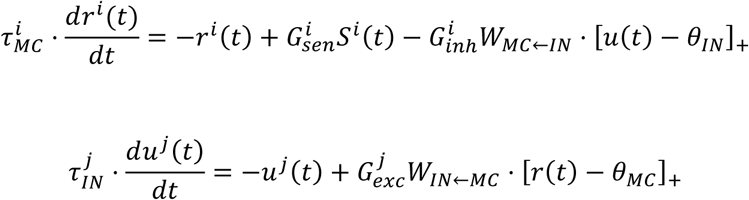

where the vectors r *θ*_*MC*_ and *θ*_*IN*_ are firing thresholds, W_MC←IN_ and W_IN←MC_ correspond to the reconstructed IN-to-MC and MC-to-IN connectivity weight matrices, respectively, and the vectors *r*(*t*) and *u*(*t*) represent the firing rates of the MC and IN, respectively. []_+_ denotes half-wave rectification:

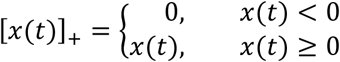

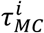 and 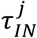 are the time constants for the individual MCs and INs, respectively. 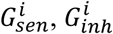 and 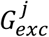 are the individual scaling factor for sensory, inhibitory and excitatory input, respectively. To account for the natural variability in biological systems, the parameter values for each of the cells in each of the individual simulation runs were drawn from a Gaussian distribution with a standard deviation of 1% of the distribution mean. The distribution means of the different parameters were:

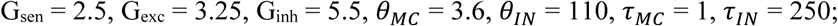

The time course of sensory input *S*^*i*^(*t*) was modelled as difference of exponentials as described previously^31^:

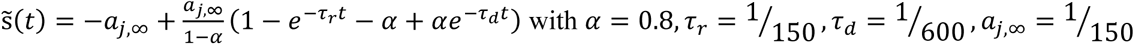

To model *S*_*i*_(*t*), the individual sensory input of MC *i*, we used its experimentally measured activity â_*i*_ during *t_1_* and modulated the time course according to 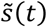:

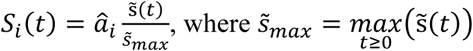

The differential equations were solved in MATLAB with a fixed step size of 1 millisecond using a first degree Newton-Cotes integration scheme or using an adaptive step size embedded Runge-Kutta-Fehlberg (4, 5) scheme. Both integration schemes lead to qualitatively very similar results, and therefore the former method was used for simplicity for the simulated data shown here.

In an iterative, semi-automated parameter search, we identified a suitable parameter range that fulfilled the following criteria:

1. The peak firing rates of individual neurons does not exceed a physiologically realistic range (< 200 Hz).
2. The strength of inhibition is appropriate to reproduce the time course of the average population activity, correlation and variance.
3. The activity, correlation contribution and variance contribution of individual MCs at *t_1_* and *t_2_* is in good correspondence to experimental measurements.

Parameters for which these criteria were fulfilled were found by parameter variations in pilot studies. Results were usually robust against variations of each parameter by ±50% around the values reported above.

### Analysis of triplet motifs

Occurrences of disynaptic MC-IN-MC motifs were counted after binarizing connections. We enumerated all neuron triplet combinations in the reconstructed wiring diagram and tested for graph isomorphism against all 4 disynaptic motif types. The obtained motif counts were compared against a reference model where the forward and backward connectivity of the MCs were permuted independently while maintaining the node count and edge density (n = 10 000 permutations). The z-scores and p-values were obtained by computing the mean and standard deviation of each motif type in the permuted networks.

To compare the motif frequency as a function of the pairwise tuning similarity, we divided the MC pairs into two groups, one with similar tuning (r_signal_ > 0.5) and one with dissimilar tuning (r_signal_ ≤ 0.5) and counted the occurrences of MC-IN-MC motifs in each group. We then compared the motif counts against a reference model where we permuted the pairwise tuning similarity between MCs and regrouped them by tuning similarity (r_signal_ > 0.5 versus r_signal_ ≤ 0.5) while maintaining the same network topology (n = 10 000 permutations). The z-scores and p-values were then obtained by computing the mean and standard deviation of each motif type in the permuted groups (Fig 4d).

### Additional analyses

The contribution of individual MCs to the Pearson correlation coefficient

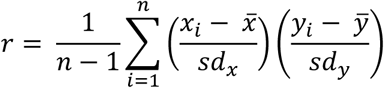

between population activity patterns was calculated by determining the summand 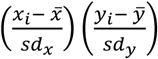 for each MC. Similarly, the contribution of individual MCs to the variance

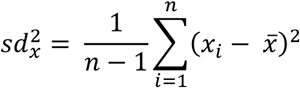

of the population activity patterns was calculated by determining the summand 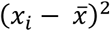 for each MC. Here, x_i_ and y_i_ are responses of MCs to odors x and y, sd_x_ and sd_y_ are the standard deviations of population responses to odors x and y, and n is the total number of MCs in the population.

Statistical significance was tested using a non-parametric Mann-Whitney U test unless noted otherwise.

